# PepCleav: a classification method for evaluating the cleavability of peptide fragments presented by MHC class I

**DOI:** 10.1101/2024.07.19.604254

**Authors:** Dawei Jiang, Wenxin Zhu, Binbin Xi, Hongli Du

## Abstract

Neoantigens, due to their tumor specificity and lack of central tolerance, are promising targets for cancer immunotherapy. Evaluating the cleavability of peptide fragments generated by proteasomes and aminopeptidases is crucial for neoantigen identification, but tools for this purpose are scarce. To address this, we developed PepCleav, a method for evaluating peptide cleavability by classifying the amino acids at the proteasome and aminopeptidase recognition sites of the C- and N-termini. By integrating C- and N-terminal cleavability, PepCleav accurately predicted the overall cleavability of peptide fragments in test datasets, correctly identifying 84-89% of cleavable peptides and 63-92% of non-cleavable peptides. We also found that highly cleavable peptides have a higher likelihood of being effective neoantigens, highlighting PepCleav’s potential to improve neoantigen identification. PepCleav’s source code is publicly available at https://github.com/Dulab2020/PepCleav.

## 1 Introduction

Neoantigens, derived from somatic mutations in cancer, can elicit anti-tumor immune responses when presented by the Major Histocompatibility Complex (MHC) to T cells. Their tumor-specific nature and ability to evade immune tolerance make neoantigens promising targets for immunotherapy (1). However, the clinical application of neoantigens has been hampered by low identification accuracy (2). A key challenge in neoantigen identification is predicting peptide fragment cleavability, which is crucial for determining whether these fragments can be generated by proteasomes and aminopeptidases and subsequently presented on the cell surface (3). Unfortunately, effective predictive tools are scarce. NetChop, one of the few available tools for predicting peptide C-terminal cleavability, uses a neural network-based method (4, 5). Despite its widespread use (6, 7), the effectiveness of NetChop remains uncertain (8). Moreover, there are no methods available for estimating the cleavability of the N-terminus, further complicating the assessment of the overall cleavability of peptide fragments.

The cleavability of peptides may be influenced by the types of amino acids at terminal enzymatic recognition sites. At the C-terminus, the proteasome subunits β1, β2, and β5 can cleave acidic, basic, and hydrophobic amino acids, respectively (9, 10, 11). At the N-terminus, aminopeptidase ERAP1 prefers to cleave hydrophobic or aliphatic amino acids (12), while ERAP2 favors acidic amino acids (13). Inspired by these observations, we conducted a statistical analysis of peptide terminals in the database, identifying the key sites for enzymatic recognition and examining the relationship between amino acids types at these sites and their cleavability. Based on these results, we have developed PepCleav, a classification method for evaluating the cleavability of peptide C- and N-termini, as well as the overall cleavability of peptides. In test datasets, PepCleav demonstrated satisfactory performance.

## 2 Methods

### 2.1 Collections and processing of MHC-I peptide ligands data

We collected all available peptide ligands from the IEDB (14) and selected those derived from cellular experiments and verified to bind to MHC I molecules (Fig. 1). Subsequently, we evaluated the peptide library saturation for each protein. The saturation is defined as:

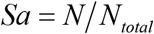

where *N*_*total*_ is the total count of de-duplicated sites detected at the C-terminus or N-terminus, and *N* is the count of sites with a coverage of two or more.

**Figure 1.**
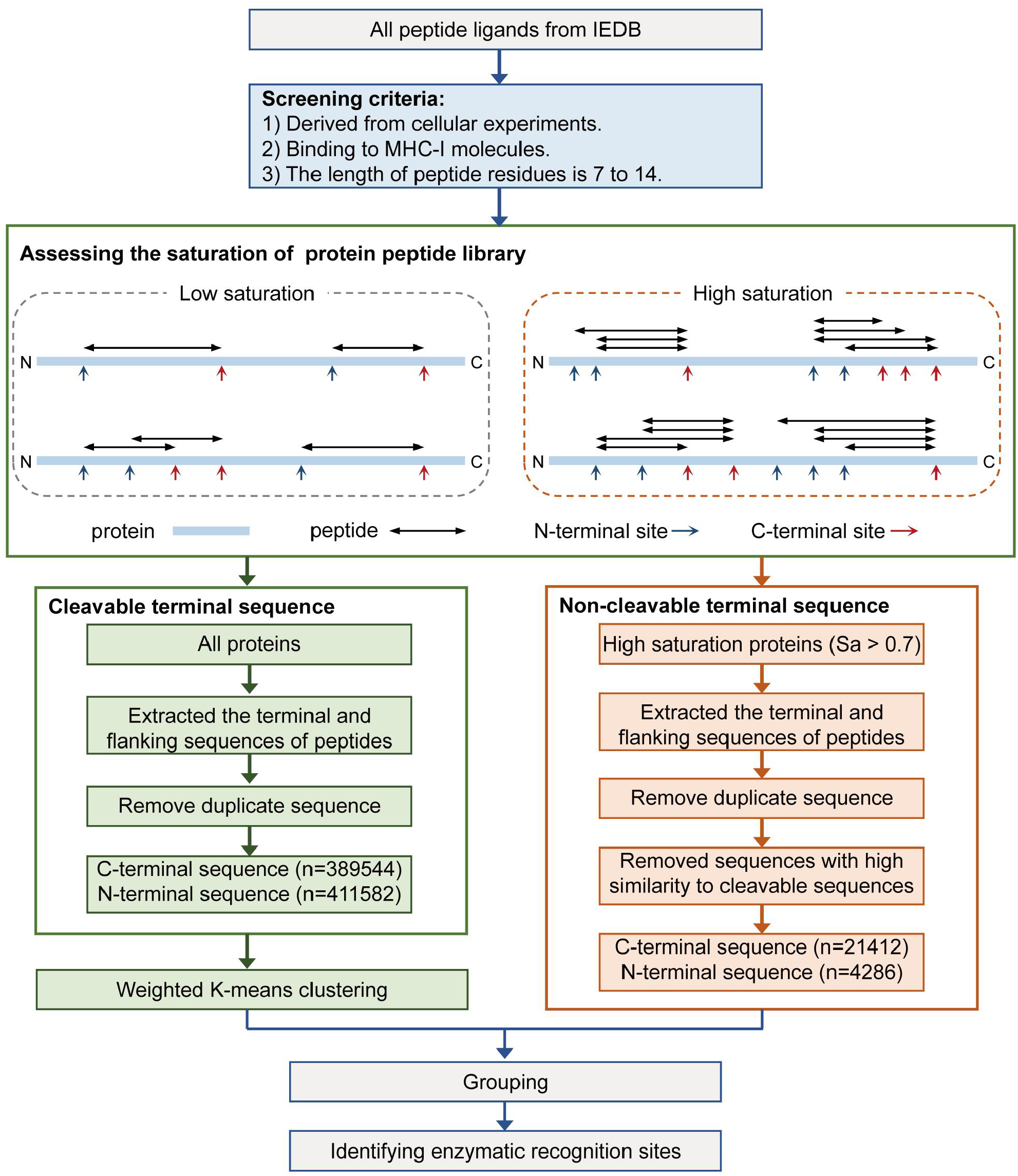
Workflow of the processing of MHC Class I ligand data. The workflow include: screening peptides from the IEDB; calculating the saturation of peptide libraries for each protein; generating cleavable sequences form all available peptide; generating non-cleavable sequences from proteins with high peptide library saturation; clustering and statistics of terminal sequences; identifying the terminal enzymatic recognition sites.

To obtain the cleavable terminal sites, we extracted the terminal and flanking sequences of these peptides from all proteins (Fig. 2A). In total, we collected 389,544 cleavable C-terminal sites and 411,582 cleavable N-terminal sites. Then we generated non-cleavable sites from the region lacking peptide coverage within proteins (Fig. 1). To avoid including erroneous samples, we meticulously selected proteins exhibiting high peptide library saturation. Empirically, proteins with *Sa* > 0.7 were retained. We then eliminated sites displaying high similarity to cleavable sites. The similarity is determined by calculating the Euclidean distance between the query sites and the cluster center of the cleavable sites, as described in methods below. Ultimately, we collected 21,412 non-cleavable C-terminal sites and 4,286 non-cleavable N-terminal sites.

**Figure 2.**
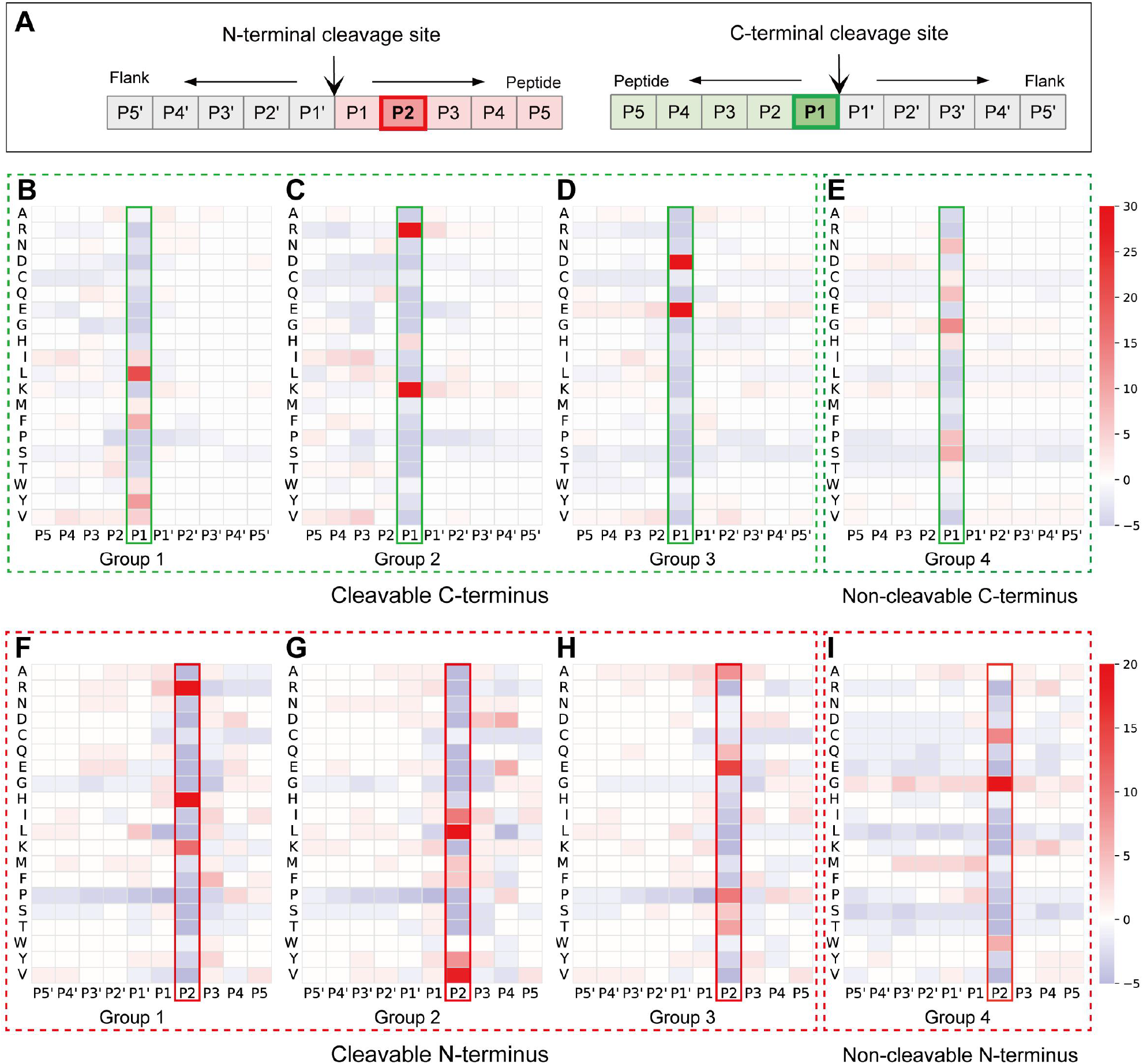
Visualization of the clustering results of terminal sites. **A**. Schematic of peptide terminal sites. **B-D**. Cleavable C-terminal sites. Group 1, Group 2 and Group 3 are three groups obtained from clustering. **E**. Non-cleavable C-terminal sites. **F-H**. Cleavable N-terminal sites. **E**. Non-cleavable N-terminal sites. The different colors represent the degree of deviation between the amino acid frequencies in each group and the background frequencies.

### 2.2 Weighted K-means clustering

To identify the terminal enzymatic recognition sites, we encoded the terminal sequences using physicochemical properties (Table S1) and then applied weighted K-means method for clustering. The core concept of weighted K-means is calculating the distance between each data point and the cluster center by multiplying a weight score. In this study, the weight score (*W*) consists of two components: *W*_*α*_ and *W*_*β*_,

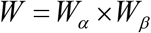

*W*_*α*_ is related to the entropy of each site. For each site *i* ∈ {P1, P2…P5, P1’, P2’…P5’}, the *W*_*αi*_ is calculated as:

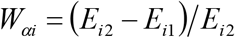

where *E*_*i1*_ is the information entropy of sequences within a cluster (cluster-sequences), and *E*_*i2*_ is the information entropy of sequences randomly generated from human proteome (random-sequences). The information entropy is calculated as:

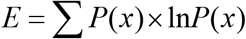

where *P*(*x*) represents the frequency of the amino acids *x* at site *i*.

*W*_*β*_ is related to physicochemical properties of amino acid. For each site *i* ∈ {P1, P2…P5, P1’, P2’…P5’} and each property *j* ∈ {Polarity, hydrophilicity, hydrophobicity, charge, volume, ……}(Table S1), we calculated the mean value of cluster-sequences (*M*_*ij1*_) and random-sequences (*M*_*ij2*_). Then *W*_β*ij*_ is calculated as:

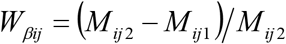

Empirically, the cleavable C-terminal and N-terminal sites were clustered into three groups (Fig. 2). The non-cleavable terminal sites groups were generated from proteins with high saturation (Fig. 1).

### 2.3 Calculation of cleavability

In order to quantify the cleavability of different types of terminus, we calculated the cleavability scores for each site in high saturation proteins. The cleavability scores is calculated as:

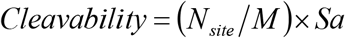

where *N*_*site*_ is the number of peptide terminus observed at each site. *Sa* is the saturation of peptide library for each protein. *M* is the mean cleavable level of each site, it is calculated as:

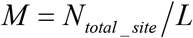

where *N*_*total_site*_ is the total number of peptide terminus observed at all site. *L* is the residue length of protein.

### 2.4 Estimation method for the cleavability of peptide

We developed PepCleav, a classification method to evaluate the cleavability of peptide fragments. It begins by categorizing termini into three classes: *Low, Medium*, and *High*, according to the median cleavability scores of amino acid at the enzymatic recognition sites (Fig. 3A-B, Table S2). Specifically, for the C-terminal P1 site, the thresholds are: scores < 1.20 (*Low*), 1.20 ⩽ scores ⩽ 1.80 (*Medium*), and scores > 1.80 (*High*). Similarly, for the N-terminal P2 site, the thresholds are: scores < 1.45 (*Low*), 1.45 ⩽ scores ⩽ 1.65 (*Medium*), and scores > 1.65 (*High*). When setting the saturation (*Sa*) threshold, we strive to maintain consistency in the physicochemical properties of amino acids within different groups. Subsequently, we classify the overall peptide cleavability into five levels, ranging from high to low, by combining the cleavability assessments of both C-terminus and N-terminus. This evaluation approach is detailed in Fig. 3C. We have developed a Python program to implement PepCleav, with the source code publicly available at https://github.com/Dulab2020/PepCleav.

**Figure 3.**
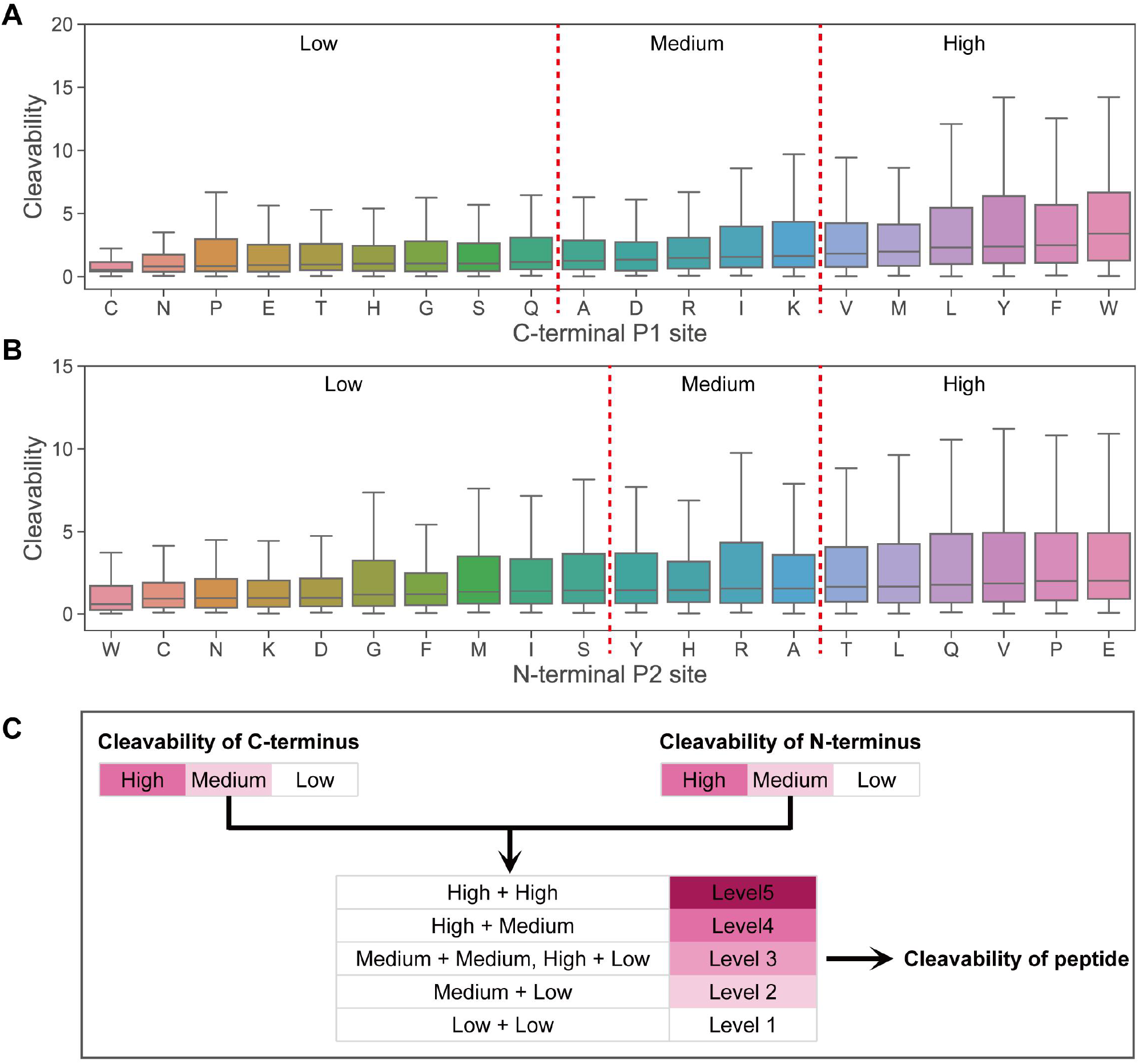
The statistics of terminal cleavability and schematic of the cleavability evaluation method. **A**. The cleavability of different amino acids at C-terminal P1 site. **B**. The cleavability of different amino acids at N-terminal P2 site. All amino acids at the key site are categorized into three groups: *Low, Medium*, and *High*. **C**. The evaluation method for the cleavability of peptide.

### 2.5 Test datasets

We have designed three distinct datasets to evaluate PepCleav’s performance (Table S3): 1) MHC-I peptide ligands dataset. This dataset contains 10,000 peptides, with an approximately 1:1 ratio of cleavable and non-cleavable peptides. The cleavable peptides were randomly selected from IEDB. The non-cleavable peptides were generated from high saturation proteins and subsequently filtered to exclude those containing cleavable terminus. 2) HIV proteins epitopes dataset. This dataset contains 2,047 cleavable peptides from HIV CD8+ epitopes dataset (https://www.hiv.lanl.gov), and 3,729 non-cleavable peptides. The non-cleavable peptides were generated from four HIV proteins (Env, Pol, Gag, and Nef) using the method described above. 3) Tumor neoantigen epitopes dataset. This dataset comprises 1,282 immunogenic peptides and 6,476 non-immunogenic peptides (8). We utilized the immunogenic peptides to assess PepCleav’s effectiveness, while employing the entire dataset to investigate the correlation between peptide cleavability and immunogenicity.

### 2.6 Performance measure

When evaluating the performance of PepCleav, *Low* is treated as the negative, and *Medium* and *High* as the positive. For the cleavability of peptide, *Level 1* and *Level 2* are treated as the negative, and *Level 3, Level 4*, and *Level 5* as the positive. The performance of PepCleav is evaluated using sensitivity and specificity, which are defined as:

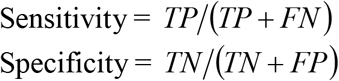

where *TP* is the number of true positives, *TN* is the number of true negatives, *FN* is the number of false negatives, and *FP* is the number of false positives.

One-sided Mann-Whitney U tests is used to measure the enrichment of immunogenic neoantigens among peptides with high cleavability level.

## 3. Results

### 3.1 Identifying the terminal enzymatic recognition sites

At the C-terminal P1 site, three cleavable clusters exhibited distinct amino acid enrichment patterns (Fig. 2B-D), potentially reflecting the specific β-subunit type of the proteasome (10). Notably, group 3, 2, and 1 may correspond to the proteolysis products of β1, β2, and β5 subunits, respectively. This correlation aligns with the β1 subunit’s caspase-like activity (post acidic amino acid), β2’s trypsin-like activity (post basic amino acid), and β5’s chymotrypsin-like activity (post hydrophobic amino acid) (11). Additionally, the presence of amino acids C, N, P, E, T, H, G, S, and Q at the P1 site may indicate reduced C-terminal cleavage propensity (Fig. 2E).

At the N-terminal P2 site, three cleavable clusters also revealed distinct amino acid signal enrichments (Fig. 2F-H). Considering that ERAP1 favors hydrophobic or aliphatic N-terminal amino acids (12) and less effectively processes G, S, and K as substrates (15), while ERAP2 prefers basic N-terminal amino acids (13), we can infer that group 2 likely represents ERAP1 proteolysis products, and group 1 corresponds to ERAP2 products. Group 3 may comprise a mixture of products from ERAP1, ERAP2, and the proteasome. The presence of amino acids C, W and G at the P2 site may indicate reduced N-terminal cleavability (Fig. 2I).

In conclusion, the C-terminal P1 site and the N-terminal P2 site are the key enzymatic recognition sites.

### 3.2 Statistical analysis of peptide terminal cleavability

We analyzed the cleavability of different amino acids at terminal enzymatic recognition sites. At the C-terminal P1 site, amino acids C, N, P, G, H, T, S, and Q exhibit relatively low cleavability, aligning with the C-terminal characteristic signal of group 4. Conversely, amino acids V, M, F, Y, L, and W demonstrate high cleavability, corresponding to group 1 (Fig. 3A). At the N-terminal P2 site, no clear correlation exists between clustering groups and cleavability (Fig. 3B). Based on these terminal cleavability statistics, we developed PepCleav to evaluate the cleavability of peptide termini and fragments (see Materials and methods).

### 3.3 Evaluating the cleavability of terminus

PepCleav demonstrated superior predictive performance for C-terminal cleavability across all test sets compared to NetChop (Table 1). Despite its weakest performance in the HIV proteins epitopes dataset, PepCleav still achieved 85.9% sensitivity and 89.4% specificity.

**Table 1.** Predictive performance for the C-terminal cleavability.

| Test datasets | Model | Sensibility | Specificity |
| --- | --- | --- | --- |
| MHC-I peptide ligands | PepCleav | 91.42% | 93.21% |
|  | NetChop3.1 | 86.09% | 62.82% |
| HIV proteins epitopes | PepCleav | 85.93% | 89.84% |
|  | NetChop3.1 | 77.87% | 92.49% |
| Cancer neoantigen epitopes | PepCleav | 98.05% | — |
|  | NetChop3.1 | 88.38% | — |

The predictive performance for N-terminal cleavability was comparatively lower. In the MHC-I peptide ligands dataset, PepCleav achieved 76.0% sensitivity and 82.2% specificity, while in the HIV proteins epitopes dataset, it reached 72.3% sensitivity and 51.5% specificity (Table 2).

**Table 2.**
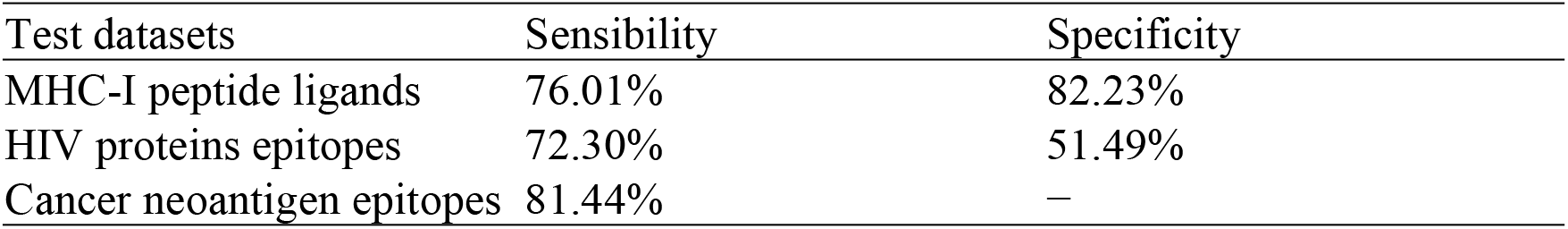
Predictive performance for the N-terminal cleavability.

### 3.4 Evaluating the cleavability of peptide fragments

PepCleav also exhibited satisfactory predictive performance for peptide fragment cleavability. In the MHC-I peptide ligands dataset, PepCleav achieved 88.9% sensitivity and 91.9% specificity. In the HIV proteins epitopes dataset, it attained 83.6% sensitivity and 62.8% specificity (Table 3).

**Table 3.**
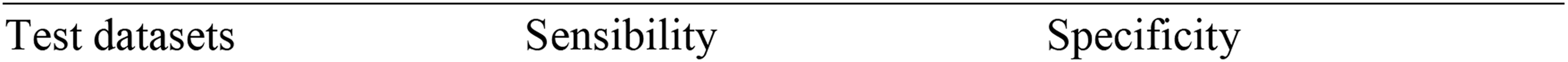
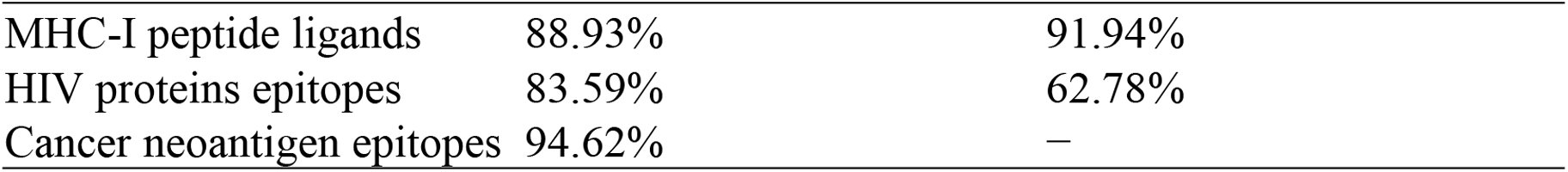
Predictive performance for the cleavability of peptide fragments.

| Test datasets | Sensibility | Specificity |
| --- | --- | --- |
| MHC-I peptide ligands | 88.93% | 91.94% |
| HIV proteins epitopes | 83.59% | 62.78% |
| Cancer neoantigen epitopes | 94.62% | — |

To further enhance the clinical credibility of our method, we investigated the potential correlation between peptide cleavability and tumor neoantigen immunogenicity. Our findings indicate that peptides with high cleavability levels are more likely to be immunogenic neoantigens (Fig. 4), a statistically significant result (*p* < 0.001) as determined by a Mann-Whitney U test.

**Figure 4.**
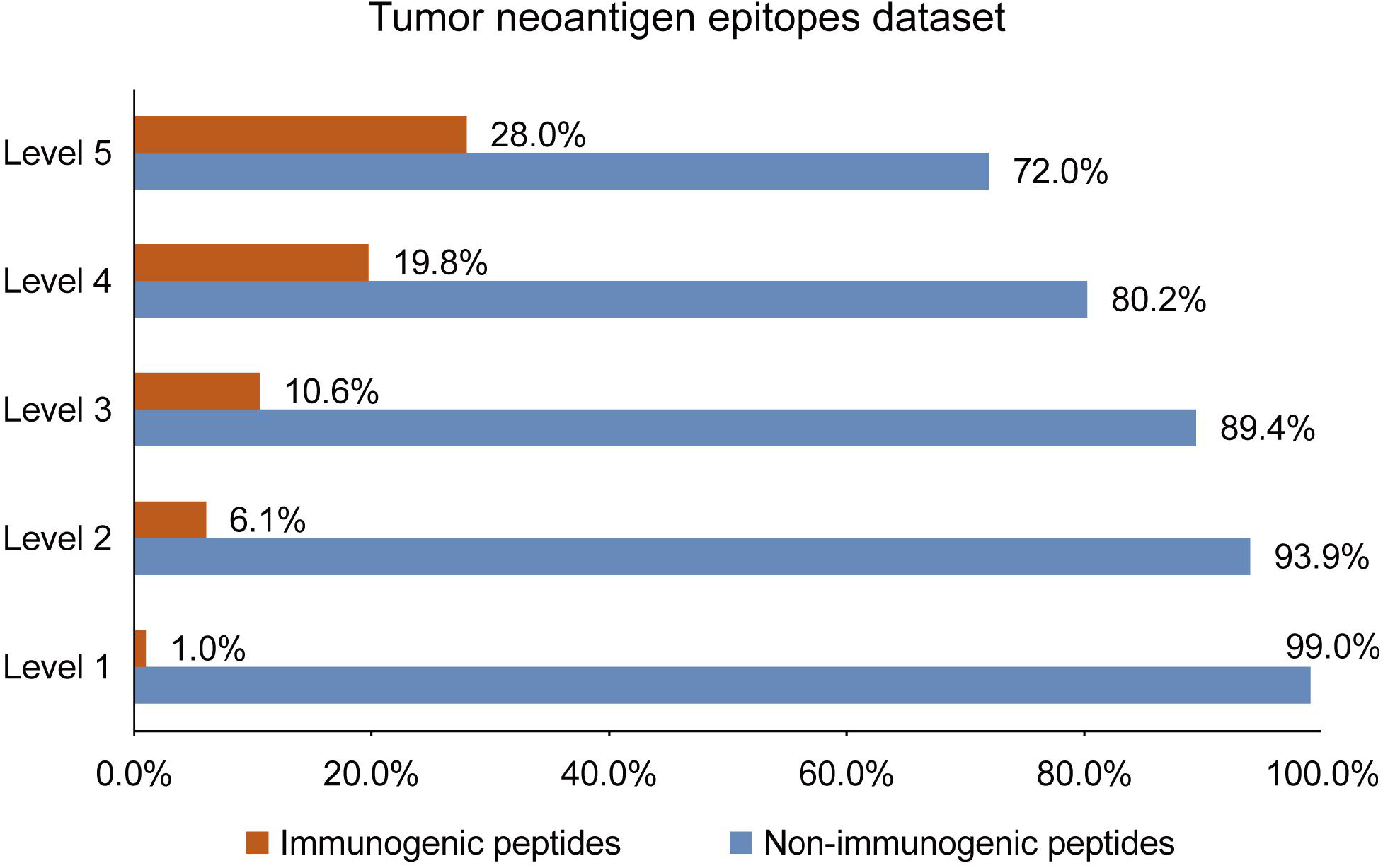
Enrichment of immunogenic neoantigens among peptides with different levels of cleavability. Peptides with higher cleavability levels are more likely to be immunogenic neoantigens.

## 4 Discussion

We provide PepCleav, a highly feasible method for predicting the cleavability of peptide presented by MHC class I. Unlike neural network models such as NetChop (5), PepCleav employs a classification approach based on statistical analysis of amino acids at terminal enzymatic recognition sites. In three test datasets, PepCleav has demonstrated robust predictive capability for peptide cleavability. Moreover, we have confirmed a correlation between high peptide cleavability and increased likelihood of immunogenic neoantigens.

Nevertheless, we acknowledge certain limitations of our method. Firstly, our study primarily focuses on the influence of key active sites on cleavability. While supported by clustering results, this approach still lacks definitive experimental validation. Secondly, cleavability is not the sole determinant in identifying immunogenic neoantigens (2). Additional factors, such as antigen presentation and antigen foreignness (16, 17), should be integrated into comprehensive neoantigen prediction analyses.

In conclusion, our study suggests that PepCleav shows promise in enhancing peptide cleavability prediction and serves as a valuable tool for identifying immunogenic neoantigens. Further research and refinement could potentially expand its applications in immunology and cancer research.

## 8 Supplementary Material

Table S1. Physicochemical properties of amino acids

Table S2. Stat of cleavability for different types of terminus

Table S3. Stat of test datasets

## Notes

### Competing Interest Statement

The authors have declared no competing interest.

